# FAβ-gal: an automated fluorescence-based quantification of the senescence-associated beta-galactosidase X-gal assay

**DOI:** 10.64898/2026.02.27.707795

**Authors:** Antonio G. Tartiere, David Roiz-Valle, Yaiza Español, Gabriel Bretones, José M. P. Freije, Alejandro P. Ugalde

## Abstract

Cellular senescence plays a pivotal role in aging and cancer, two major biomedical and socioeconomic challenges of our time. Therefore, its study has become crucial for the design of interventions based on its manipulation. In this sense, researchers have developed a wide variety of techniques to detect and quantify cellular senescence. Among them, the most popular is the original Senescence-Associated β-galactosidase (SA-β-gal) colorimetric assay, based on the use of the chromogenic substrate X-gal. This compound is cleaved by β-galactosidase, producing an insoluble, blue precipitate of 5,5’-dibromo-4,4’-dichloro-indigo (commonly referred to as indigo). While this method remains the gold standard senescence assay, its quantification remains challenging due to the color-based readout. In this work, we describe a method, which we have named FAβ-gal (Fluorescence Analysis of β-galactosidase), that exploits the far-red fluorescence of the β-gal product indigo and allows the quantification of SA-β-gal activity under any conventional wide-field fluorescence microscopy using the original X-gal assay. In addition, we developed workflows and software applications that standardize SA-β-gal quantification in a semiautomatic and unbiased manner. We demonstrate that FAβ-gal measurements present a strong linear correlation with the percentage of senescent cells and show high sensitivity. Moreover, we show that this method is also applicable to tissue sections, underscoring the versatility of our approach. Therefore, FAβ-gal could be easily introduced into the routine of laboratories already using the original colorimetric assay, enhancing the accuracy, sensitivity and reproducibility of senescence detection.

## Introduction

More than sixty years ago, it was discovered that human diploid fibroblasts could only overcome a finite number of cell divisions before entering a state of permanent proliferation arrest. This state was named cellular senescence and was caused by the telomere attrition that occurs with each cell division, leading to the shortening of these protective structures^1^. Nowadays it is known that cellular senescence can be triggered by different stressors and not only by telomere shortening. In this sense, several forms of stress such as DNA damage, oxidative stress or oncogene activation have been shown to induce different types of cellular senescence that share a common phenotype characterized by an irreversible proliferation arrest and an intense Senescence Associated Secretory Phenotype (SASP)^2^. On the other hand, it has been demonstrated that cellular senescence plays a pivotal role in physiological homeostasis, contributing to tissue development, wound healing and tumor supression^3^. However, the clearance mechanisms that eliminate these cells after they have accomplished their function decline with age, leading to their accumulation. In this sense, this increase in senescent cells is considered a hallmark of aging as it contributes to this process by disrupting tissue homeostasis and promoting chronic inflammation and gradual fibrosis^4^. Moreover, as this low-grade inflammation and tissue fibrosis contribute to tumorigenesis, senescent cells have also been proposed as a hallmark of cancer^5^. Hence, cellular senescence exerts an ambivalent role in cancer development. In this sense, its induction after damage prevents the transformation of damaged cells into neoplastic cells. However, if these cells are not removed by the immune system, as it occurs with age, their accumulation creates a profibrotic and proinflammatory environment that stimulates tumor progression^6^.

As cellular senescence plays a key role in aging and cancer, some of the biggest sanitary and socioeconomic challenges of the moment, its understanding has become crucial. In this sense, several studies have demonstrated that the elimination of senescent cells with targeted drugs, known as senolytics, increases healthspan and lifespan in mouse models of different diseases and in aged mice^7,8^. In addition, multiple works have shown that interventions designed to induce senescence of tumor cells can exert an antineoplastic activity through the immunostimulant and proangiogenic effect of SASP^9,10^. Moreover, some studies have demonstrated that the use of these therapies followed by a treatment with senolytics also impairs tumor growth^11^. Hence, senolytics and pro-senescence therapies have emerged as promising approaches for aging and cancer management, and their development requires methods to detect and quantify cellular senescence.

In this sense, the first method to detect senescent cells, which still remains the gold standard, was described in 1995 and is based on the beta-galactosidase (β-gal) activity that senescent cells display at pH 6. This activity, referred to as Senescence-Associated β-galactosidase (SA-β-gal), marks senescent cells and is detected using the chromogenic substrate 5-bromo-4-chloro-3-indolyl β-D-galactopyranoside (X-gal). Cleavage of this compound by β-gal yields a blue, insoluble 5,5’-dibromo-4,4’-dichloro-indigo (indigo) precipitate^12^. Thus, senescent cells can be identified using this colorimetric assay, which is simple, economical and fast. However, the need for more sensitive and quantitative methods led to the development of new techniques that involve fluorescence assays. In this regard, some of these methods maintain the biological basis of the colorimetric assay but use fluorogenic substrates of β-gal such as 5-dodecanoylaminofluorescein di-β-D-galactopyranoside (C_12_FDG), enabling a more sensitive and quantitative measure of senescence levels^13^. On the other hand, researchers have developed new methods based on other biological characteristics of senescent cells, such as the expression of certain biomarkers (e.g. *p16* or *p21*)^14^, their intrinsic autofluorescence^15^ or even the changes in their rRNA^16^. Nevertheless, the greater complexity, time consumption and cost of these techniques compared to the classical SA-β-gal assay has hampered their usage by the scientific community.

Hence, as mentioned above, the colorimetric SA-β-gal assay remains the gold standard for monitoring senescence, but it possesses important quantification and sensitivity limitations due to the color-based readout. On one hand, conventional SA-β-gal assays require trans-illumination, which suffers from uneven field illumination, especially in multi-well vessels due to the meniscus effect of aqueous solutions^17^. On the other hand, it requires a color camera, which has lower sensitivity than monochrome counterparts and is not always found in fluorescence microscopy setups, thereby limiting the simultaneous detection of fluorescence-based readouts (such as DAPI or Hoechst counterstain). Moreover, there is no consensus method to analyze the results of this assay. In this sense, there are a wide variety of approaches, all of them with their own advantages and limitations, such as counting manually the number of positive and negative cells of a few random fields^13^, quantifying the color intensity of the photographed cells^18^ or using deconvolution algorithms to isolate the indigo signal from the counterstain and background^19,20^. However, these strategies often require manual counting or selection of cells and can be greatly affected by the differences on illumination in bright field images, as well as by possible biases linked to the direct intervention of the researcher. These issues make these approaches time-consuming and limit their reproducibility.

Therefore, in this work we propose a new method that joins the simplicity, low cost and fast execution of the classical assay, with the quantitative character and sensitivity of fluorescence techniques, together with an unbiased and semiautomatic execution. This new approach, which we have named FAβ-gal (Fluorescence Analysis of β-gal), is based on the inherent fluorescence of the X-gal product described by Levitsky et al. in the context of LacZ reporter^21,22^, along with a deep-learning-powered count of the nuclei^23^ and a semiautomatic analysis of the images.

## Results

### Far-red fluorescence enables efficient detection of the β-gal product indigo in cells

Indigo far-red fluorescence has been used to detect LacZ expression using a confocal microscope^21^. However, we wondered whether this fluorescent property could be also harnessed to quantify SA-β-gal activity using a wide-field fluorescence microscope. As mentioned before, cell autofluorescence is a very well characterized phenomenon that can interfere with fluorescence assays, especially when using senescent cells whose green fluorescence has been described^15^. Therefore, to determine whether this autofluorescence could interfere with indigo fluorescence, we compared the signal of stained and unstained senescent human primary fibroblasts GM00038 under different wavelengths (Figure 1 and Figure S1). The results showed negligible autofluorescence in the red channel (605/70 nm) and even lower in the far-red (709/100 nm), while a marked autofluorescence was observed in the green channel (525/50 nm). In contrast, stained senescent cells showed a clear fluorescence signal in the far-red channel that correlates with the blue intensity of brightfield images (Figure 1). Hence, the far-red channel setup, which matches the excitation/emission spectrum of indigo reported by Levitsky et al.^21^, constitutes an optimal region of the light spectrum to quantify the specific fluorescence signal of the indigo SA-β-gal product in senescent cells. On the other hand, we wondered whether, as it occurs in the green channel^15^, senescent cells display a greater autofluorescence than their proliferative counterparts in the far-red channel. To this end, we measured far-red autofluorescence in unstained GM00038 senescent and proliferative fibroblasts, using a threshold to eliminate background fluorescence and normalizing by the number of cells. The results showed that even if far-red autofluorescence of senescent cells is much weaker than autofluorescence on other channels, such as green or red channel, it is still higher in senescent than in proliferative cells (Figure S2).

**Figure 1.**
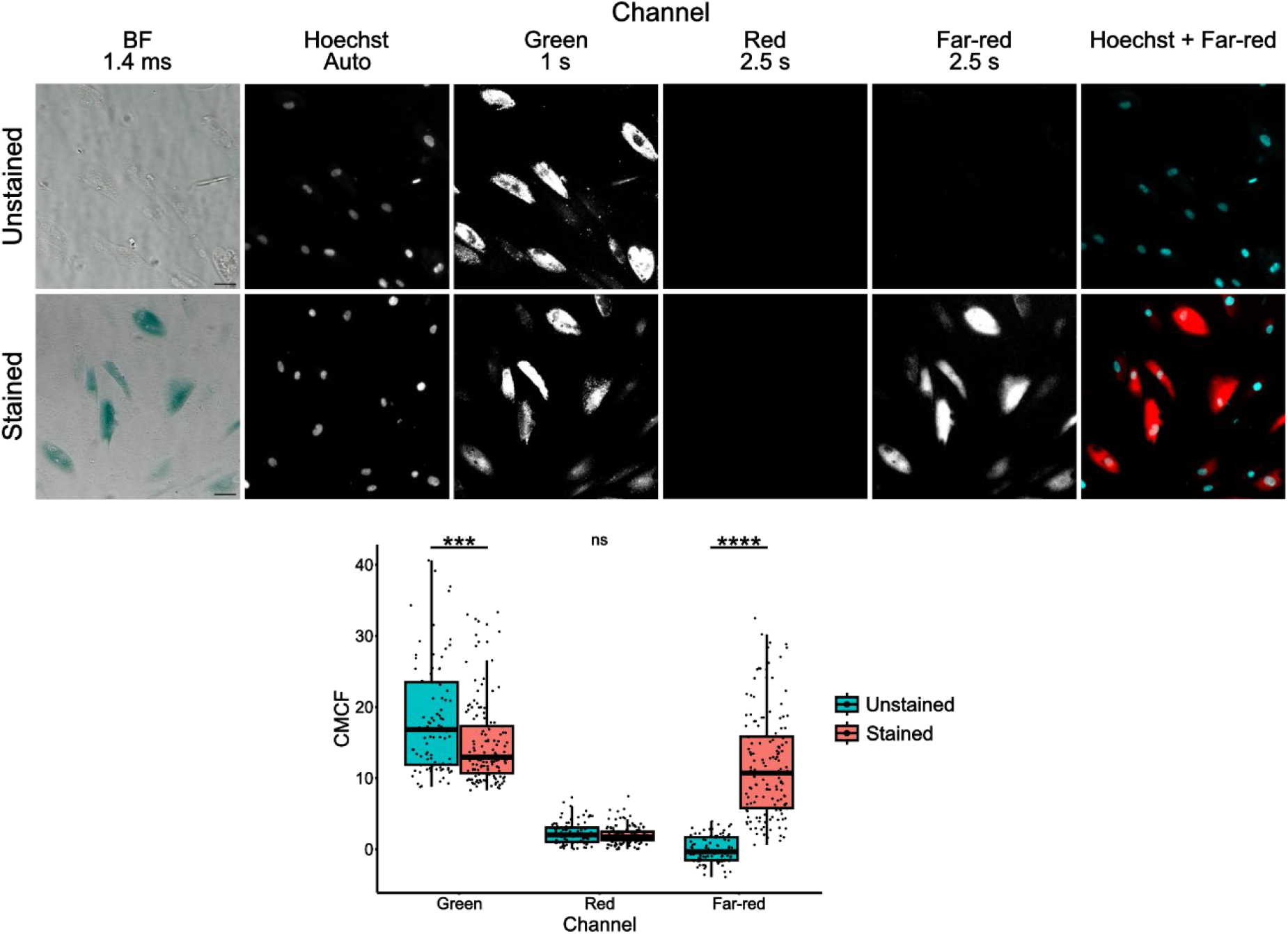
The X-gal assay product indigo can be recorded in the far-red fluorescence channel. Representative images of indigo fluorescence in X-gal SA-β-gal stained GM00038 human senescent fibroblasts compared to the autofluorescence of the same, unstained cells in the far-red, red and green channels, with the exposure times indicated. Scale bar: 50 µm. The boxplot shows Corrected Median Cell Fluorescence of X-gal stained and unstained cells (CMCF) in the green, red and far-red channels. Each point represents a cell. BF, Bright Field. *P, Wilcoxon unpaired two-sample test P-value. The statistical significance of the CMCF differences was calculated using a Wilcoxon unpaired two-sample test: *, p-value < 0.05; ***, p-value < 0.001; ****, p-value < 0.0001; ns, not significant.

These results evidenced a strong correlation between the blue precipitate intensity and its fluorescence signal, suggesting that its measurement could yield highly quantitative results. Based on these findings, we develop a method, FAβ-gal, for measuring SA-β-gal activity based on the quantification of this signal. Specifically, this method starts with a SA-β-gal staining of the cells^24^, followed by the addition of Hoechst and the semiautomatic acquisition of fluorescent images in the Hoechst and far-red channels. These images are then used as input of a pipeline that counts the nuclei using BiaPy^23^ software and quantifies the sum of the intensity values (raw integrated density) of the far-red channel positive area. This area is defined using a threshold based on the fluorescent signal of an unstained image to correct for cell autofluorescence. Then, this raw integrated density is corrected for background signal to calculate the Corrected Total Fluorescence (CTF). Finally, these data are used to compute the CTF per nucleus (CTF/nuclei) that correlates not only with the number of positive cells but also with the intensity of the stain (Figure 2).

**Figure 2.**
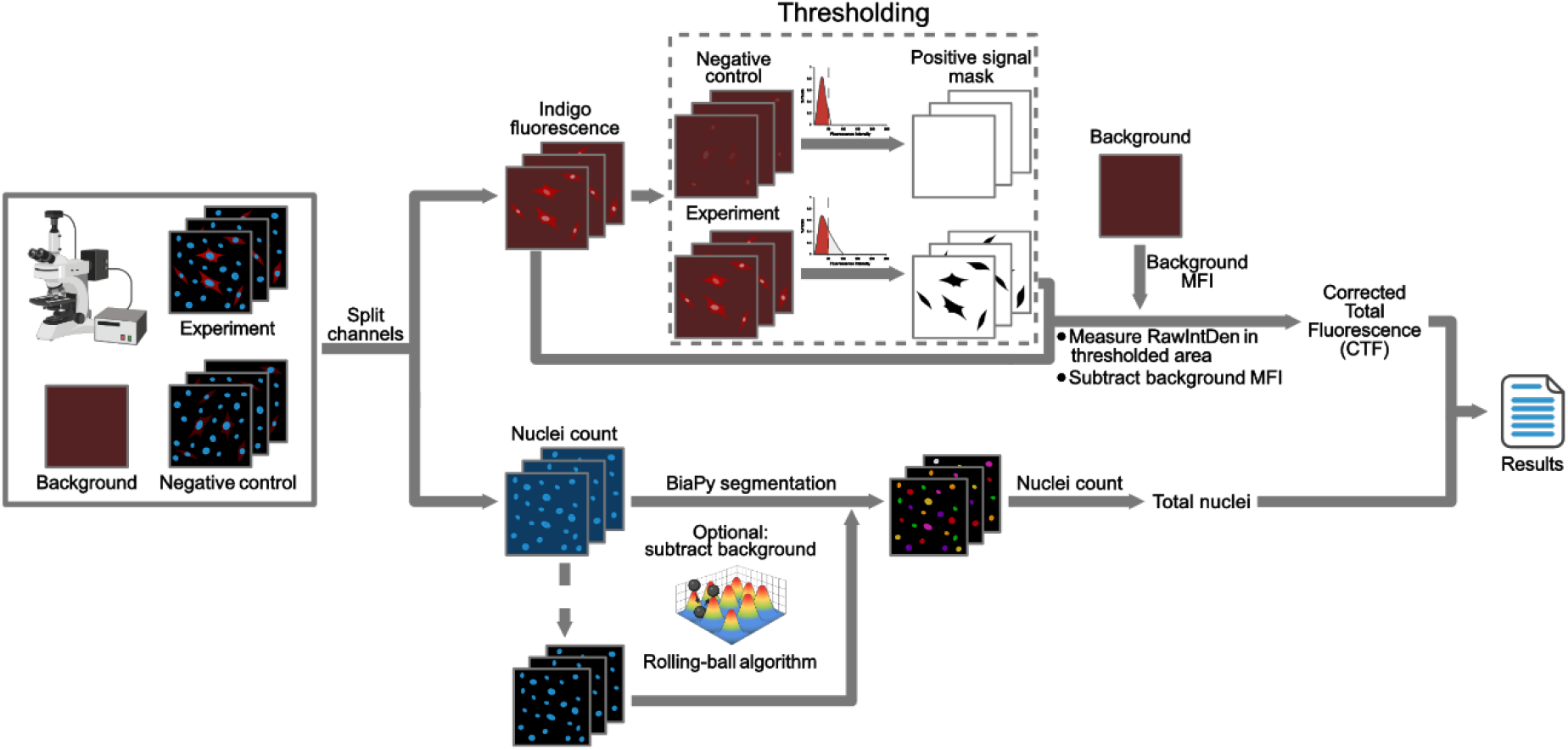
FAβ-gal analysis workflow. First, cells are subjected to a classical SA-β-gal assay and then stained with a standard nuclear dye (e.g., Hoechst). Then, fluorescence images of the nuclei and far-red channels are taken, saved as TIFF files, and manually curated to eliminate images with debris that could interfere with posterior quantification. These curated images are used as input of a pipeline that splits each channel separately. In the far-red channel, a threshold based on an unstained image (negative control) is applied to create a mask that defines the positively stained area. This mask is used to calculate the sum of the intensity values (raw integrated density, RawIntDen) of the original image only in the positively stained areas. Then, this RawIntDen is corrected for background fluorescence using the mean fluorescence intensity (MFI) of an empty image (with no cells or debris) to calculate the Corrected Total Fluorescence (CTF). On the other hand, the nuclei channel (e.g. Hoechst channel) can be optionally subjected to a background correction that uses a rolling-ball algorithm, and is used as input for BiaPy^23^. This software uses deep-learning algorithms to segmentate the nuclei, that then are counted to normalize the signal by the number of cells. Finally, these data are integrated into a results file.

### FAβ-gal is able to distinguish proliferating and senescent cells with only 3 hours of incubation

As mentioned before, the original method for quantifying SA-β-gal measured the percentage of positive cells. However, as cellular senescence is not an all-or-nothing process but a continuum, a binary approach does not reflect correctly its nature^25,26^. Hence, a method for quantifying SA-β-gal should be sensitive to reflect this continuous nature of senescence. In this sense, to simulate different levels of SA-β-gal staining and assess the sensitivity of FAβ-gal, we used GM00038 senescent and proliferative fibroblasts and increasing incubation times of X-gal assay. The results showed that FAβ-gal was able to distinguish between proliferating and senescent cells by the third hour of incubation. Moreover, it also showed a gradual increase in fluorescence signal with the time of incubation, demonstrating its ability to capture the graduality of the X-gal staining (Figure 3). In this sense, the results suggest that while the percentage of positive cells seems similar between 9 h and 24 h timepoints, the CTF/nuclei are considerably higher in the latter (Figure 3). Moreover, the comparison of the CTF/nuclei metric with other approaches such as quantifying the positive area per nuclei, showed that the relative increase in the signal with respect to the control (unstained cells) was higher in the CTF/nuclei (Figure S3). Thus, the wider range of FAβ-gal measurements allows to detect small variations in SA-β-gal signal, underscoring the sensitivity of the method.

**Figure 3.**
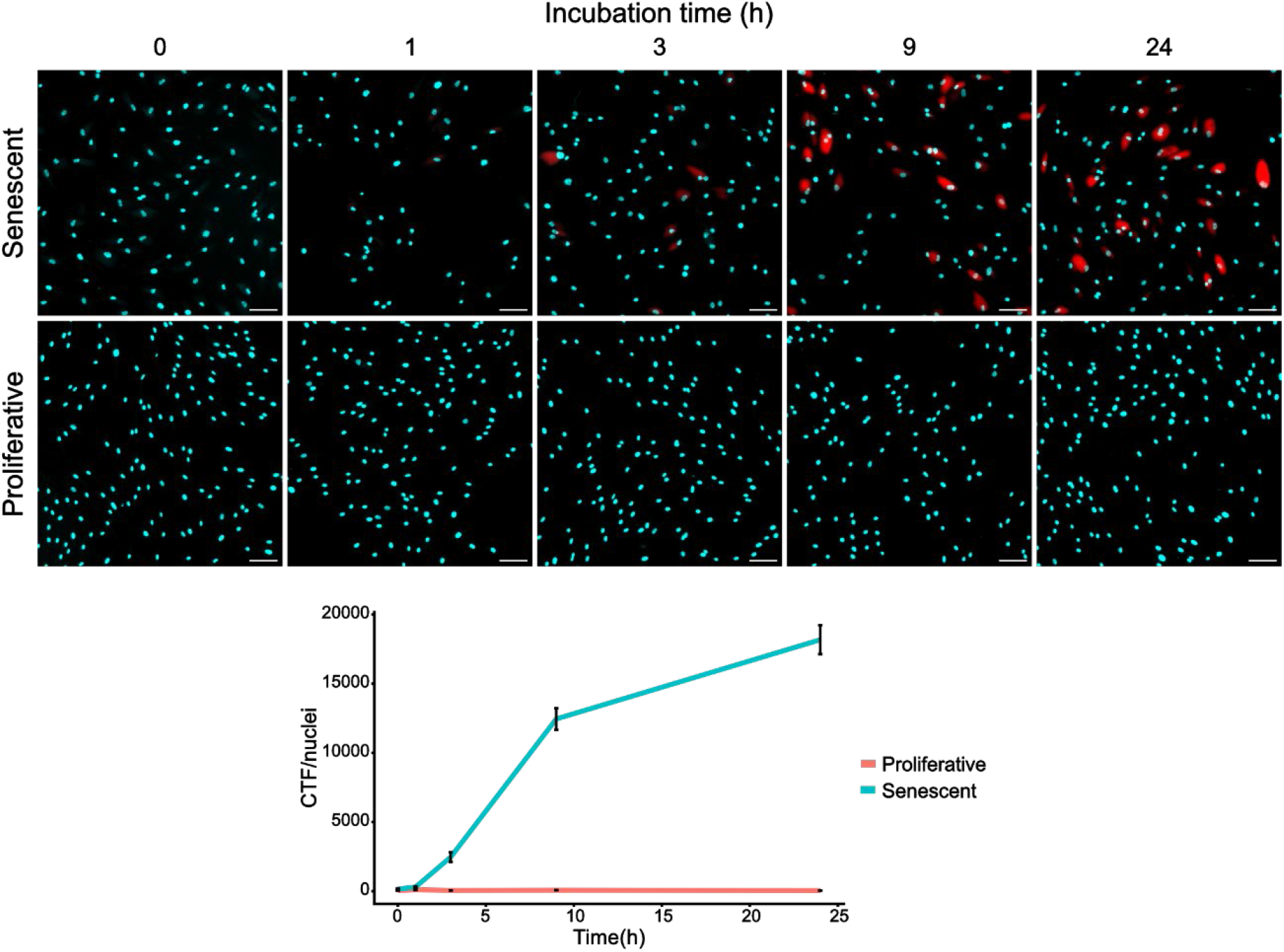
FAβ-gal captures gradual increases in indigo signal. Representative fluorescence images of GM00038 senescent and proliferative human fibroblasts stained for increasing periods of time. Blue, Hoechst channel; Red, Far-red channel. Scale bar: 100 µm. The line plot represents the evolution of the Corrected Total Fluorescence per nucleus (CTF/nuclei) of indigo with assay incubation time in proliferative and senescent GM00038 human fibroblasts. Error bars represent the standard deviation of the mean for each time point and condition.

### FAβ-gal shows a strong correlation with senescence levels

Once we have established that FAβ-gal is able to detect SA-β-gal activity with high sensitivity, we aim at testing the ability of the proposed approach to measure the percentage of senescent cells in a relevant *in vitro* culture. To achieve this goal, we mixed proliferating and replicative-senescent GM00038 fibroblast cultures in different proportions and applied our approach to evaluate the correlation of FAβ-gal measurements with the percentage of senescent cells. The results showed that FAβ-gal method exhibits a strong linear correlation (R^2^ = 0.91, p-value = 8.491x10^-10^) between the CTF/nuclei calculations and the percentage of senescent cells (Figure 4). These results underscore the high accuracy and the linear behavior of FAβ-gal method, pointing out its quantitative character and reproducibility.

**Figure 4.**
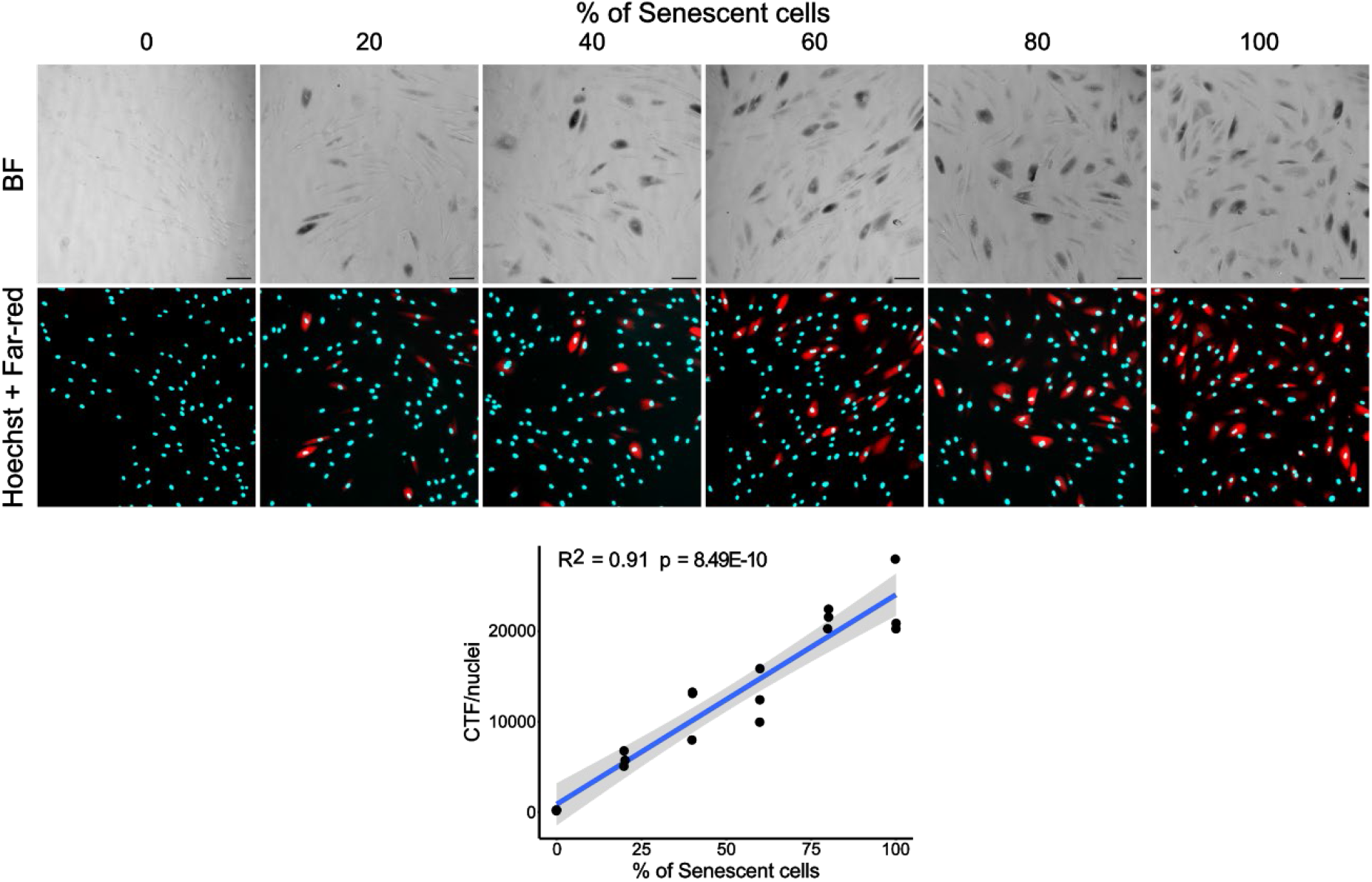
FAβ-gal shows a strong linear correlation with the percentage of senescent cells. Representative bright field and fluorescence images of combined cultures of actively proliferating and senescent GM00038 human fibroblasts. Scale bar: 100 µm. The scatter plot represents the linear relationship between the Corrected Total Fluorescence per nucleus (CTF/nuclei) of indigo and the percentage of senescent cells. Each point represents the data of a well, which is composed of several images. BF, Bright Field. *\*P*, Pearson correlation test *P*-value; R^2^, linear regression square R.

On the other hand, to evaluate the performance of the FAβ-gal method in a different cellular model we applied our approach to U2OS cancer cells treated with the topoisomerase inhibitor doxorubicin, which is known to trigger DNA damage-induced senescence^27^ Thus, we treated U2OS cells with increasing doses of doxorubicin and in two different ways, pulse and continuous, and senescence levels were quantified at different time points since the exposure using our approach. The results showed that, as expected, senescence levels increased with time after treatment in all conditions. Moreover, the senescence levels determined by our FAB-Gal method showed a clear dose-dependent increase in cells treated with a 24h pulse of doxorubicin, reaching a plateau at 50 nM. Conversely, continuous exposure to doxorubicin resulted in higher senescence levels than pulse treatments, even at lower concentrations, suggesting that sustained drug exposure drove virtually the entire U2OS cell population into senescence, whereas pulse treatment induced senescence only in a fraction of the cells (Figure 5). Altogether, these findings demonstrate that FAβ-gal is a versatile tool for quantifying different types of cellular senescence.

**Figure 5.**
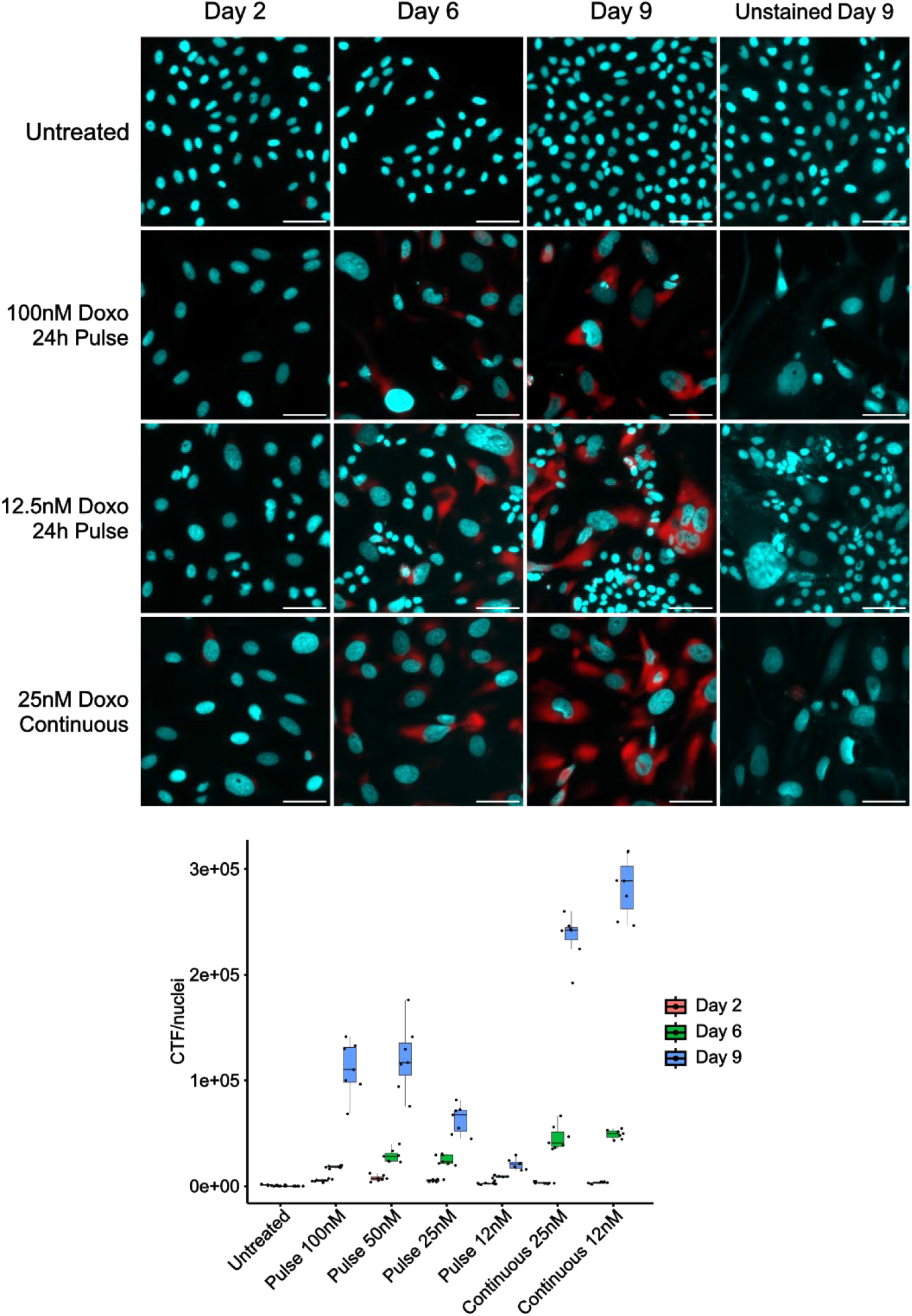
FAβ-gal is able to quantify senescence induced by doxorubicin. Representative fluorescence images of U2OS cells at different time points, treated with different doses of doxorubicin (Doxo) for 24 h or continuously. Blue, Hoechst channel; Red, Far-red channel. Scale bar: 100 µm. Untreated cells were seeded 48 h before each time point to avoid unspecific staining due to excessive confluence. The boxplot represents the Corrected Total Fluorescence per nucleus (CTF/nuclei) of indigo for each condition and time point. Each point represents the data of a well, which is composed of several images.

### FAβ-gal can be used to quantify senescence in tissue sections

One of the main advantages that has maintained the classical SA-β-gal colorimetric assay as the gold standard is that it can be used both in culture cells and tissue sections. Therefore, we assessed whether FAβ-gal could also be used in this context. To this end, we stained kidney OCT frozen sections of *Lmna*^*G609G/G609G*^ mice (LAKI), in which previous studies from our group had found increased senescence levels^28^, together with WT control samples. In order to apply Faβ-gal to tissue samples, we followed a different approach: we first selected areas from the cortex of the kidney and calculated the CTF as done with cells, but then we normalized it by the size of the area in square micrometers. Hence, in this case we computed the Corrected Total Fluorescence per area (CTF/area). The results showed that the far-red fluorescence signal is also able to capture the indigo staining specifically above the background autofluorescence in tissue sections. In line with this, FAβ-gal results evidenced a clear increase in senescence levels in the kidney of LAKI mice, as we had reported before^28^ (Figure 6). Taken together, these results support that FAβ-gal constitutes a more quantitative and sensitive approach than the classical SA-β-gal colorimetric assay in all of its applications, but maintaining its simplicity, low cost and reduced time consumption.

**Figure 6.**
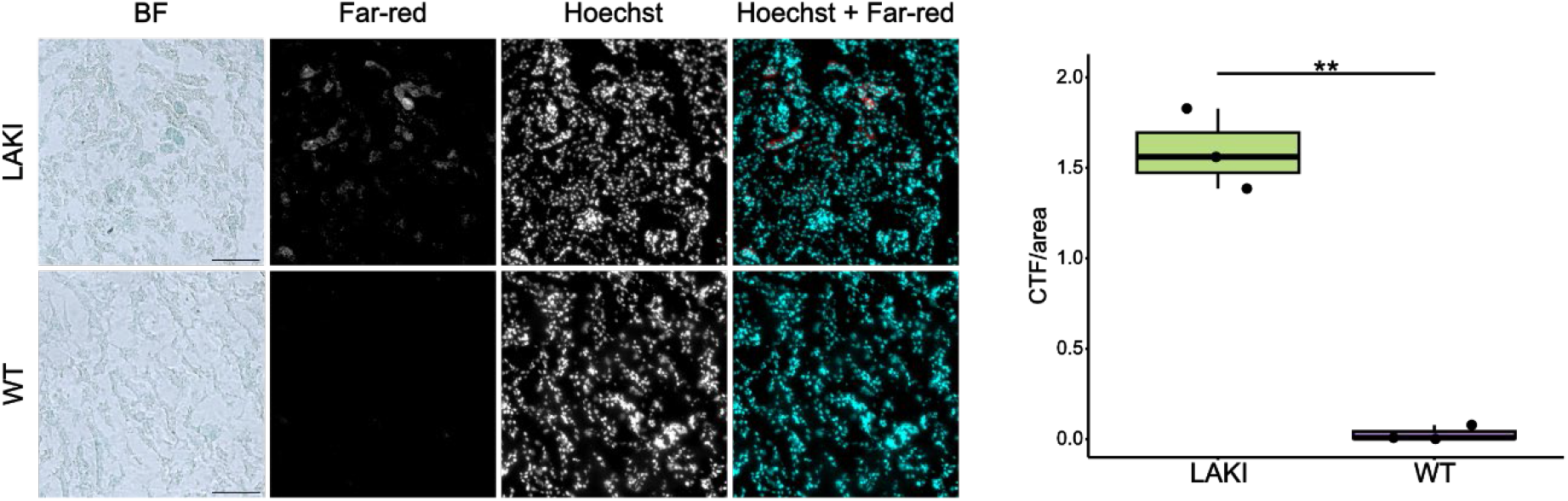
FAβ-gal is suitable for quantifying senescence in tissue sections. Representative bright field and fluorescence images of kidney cortex OCT sections from *Lmna*^*G609G/G609G*^ (LAKI) and WT mice. Scale bar: 100 µm. The boxplot represents the Corrected Total Fluorescence per area (CTF/area) of indigo in LAKI and WT mice, each point represents data of an individual. BF, Bright Field. The statistical significance of the CTF/area differences was calculated using a Student’s unpaired two-tailed t-test, with independent variances: **, p-value < 0.01.

### FAβ-gal can be used as a user-friendly application as well as a high-throughput computational pipeline

The time required to acquire computational skills has hampered the shift from manual analysis to unbiased and faster automated approaches, especially when analyzing low numbers of images. For this reason, we have developed a user-friendly R Shiny application that enables applying FAβ-gal without the need for computer skills. This application can be installed on any operating system (Windows, macOS or Linux) and allows the user to easily apply our method to a set of images. In this sense, FAβ-gal app ensures that this method can be easily incorporated into any laboratory routine (Figure 7). On the other hand, for scientists with informatics expertise and a large number of images, we have developed a Python computational pipeline. This pipeline installs all necessary dependencies and executes a customizable step-by-step analysis, allowing the users to optimize the method for their specific conditions (Figure 7). The main difference between the R Shiny app and the computational workflow is the nuclei segmentation method. In this regard, while FAβ-gal app uses a threshold-based segmentation to count the number of nuclei, the pipeline uses deep-learning software BiaPy^23^. This difference can be especially relevant when nuclear staining is not uniform, as threshold-based segmentation tends to fail in these conditions. Thus, if nuclear stain is not uniform, a fixed threshold will tend to overlook tenuously dyed nuclei and to fuse strongly dyed nearby nuclei. Finally, in a laptop with an AMD Ryzen 7 7840HS w/ Radeon 780M Graphics (3.80 GHz), 32 GB of RAM and no gpu FAβ-gal app is able to process 22 images/min and the Python computational pipeline 12 images/min. However, the speed of the computational pipeline increases substantially when a gpu is used and the background subtraction algorithm is disabled. Hence, FAβ-gal app constitutes an easy-to-use option with no additional experience needed, while FAβ-gal computational workflow is an option suitable for high-throughput analyses and is robust to differences in nuclear staining.

**Figure 7.**
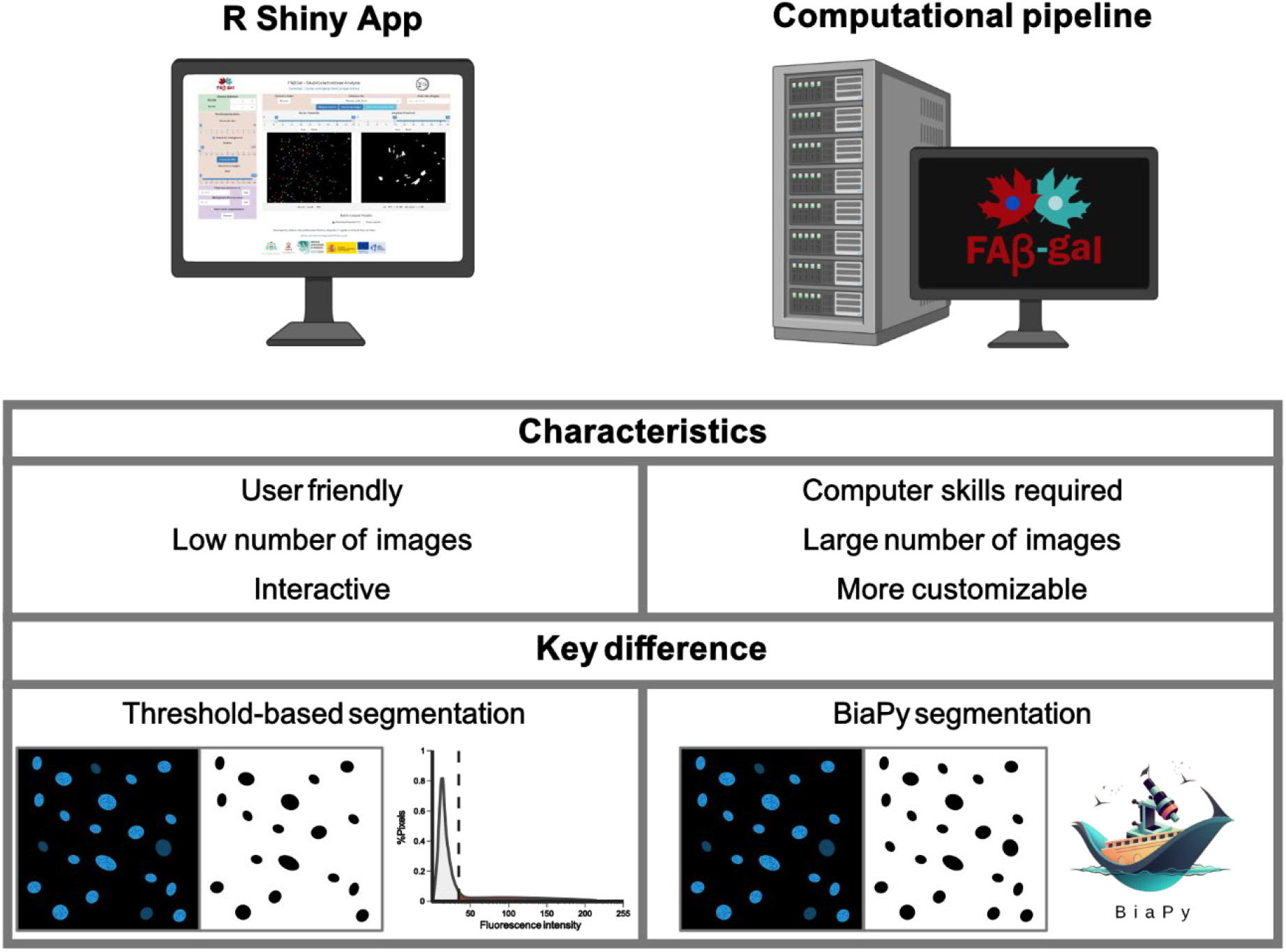
FAβ-gal quantitation software options. FAβ-gal R Shiny application constitutes an interactive and user-friendly option for analyses of low numbers of images. On the other hand, the FAβ-gal Python computational pipeline offers a faster and more customizable alternative for users with informatic expertise and large numbers of images. The main difference between these approaches lies in the segmentation step. While FAβ-gal app uses a threshold-based segmentation, the computational workflow applies BiaPy, a deep-learning segmentation algorithm more robust against uneven nuclear staining.

## Discussion

Despite the recent introduction of highly sophisticated methods for senescence quantification, the original SA-β-gal assay has remained the gold standard. This success stems from its simplicity, versatility, low cost and reduced time consumption. FAβ-gal merges all these advantages with the more quantitative and sensitive nature of fluorescence-based assays. Moreover, it can be used to detect and quantify different types of senescence, as it is based on the already proved SA-β-gal activity, both in culture cells and tissue sections. This versatility makes FAβ-gal stand out among other fluorometric methods that are usually limited to culture cells because of the strong autofluorescence that tissue sections display in the green spectrum where these techniques measure the signal^29^. In addition, this method does not require expensive fluorogenic substrates such as C_12_FDG or SPiDER-β-gal^13,29^, whose signal may be compromised by the intrinsic green autofluorescence of senescent cells. Instead, it relies exclusively on the reagents used in the original assay together with a standard nuclear fluorescent dye that does not emit in the far-red spectrum (e.g. Hoechst or DAPI). In terms of time consumption, FAβ-gal maintains the fast execution of the SA-β-gal colorimetric assay, something critical for senescence screenings and high-throughput applications.

On the other hand, our approach offers an alternative to the manual counting of the cells and their classification as positive or negative, thus accelerating the evaluation of the method and eliminating the possibility of observer-introduced bias. However, users can still apply cell counting analysis and still benefit from the more sensitive and robust detection of SA-β-gal activity and the simultaneous detection of cell nuclei of the FAβ-gal method. Moreover, the fluorescence-based nature of FAβ-gal enables advanced users to combine SA-β-gal detection with nuclei and cytoplasmic markers and use machine-learning models to segment individual cells and carry out automated measurements of SA-β-gal activity in individual cells. Furthermore, the use of fluorescence instead of the classical bright field images largely eliminates the illumination differences that appear when using small wells, such as 96-well plates, due to the meniscus effect^17^. This is critical since differences in illumination can affect posterior quantification, thus hampering the use of small wells which are fundamental for high-throughput applications. Moreover, as discussed before, the mathematical approach of this method reflects the continuous nature of cellular senescence, in contrast with those methods that calculate percentage of positive cells, and corrects the signal both by fluorescence background and autofluorescence.

However, FAβ-gal has certain limitations. In this regard, as mentioned before, autofluorescence is an intrinsic characteristic of certain culture cells and therefore, it can interfere with fluorescence measurements. Although we have demonstrated that autofluorescence in our measuring channel (far-red) is very low if compared with other channels and with indigo’s signal (Figure 1), it still reduces the sensibility of the method. Moreover, we have observed that this autofluorescence is greater in senescent than in proliferative fibroblasts (Figure S2), which could lead to an insufficient or excessive autofluorescence correction if a unique negative control is used for all conditions (something common for practicality). Another limitation of this method stems from the measurement of a precipitate fluorescence, which does not behave as a molecule in suspension. This could lead to a non-linear relationship between β-gal activity and fluorescence signal. Nevertheless, our data show that, at least under the conditions described herein, FAβ-gal shows a strong linear correlation with the percentage of senescent cells as well as a considerable sensitivity. Another critical aspect is that FAβ-gal does not provide measurements at cell level but at image level, which could difficult certain type of analyses. It should be noted that cell level measurements require additional markers for cell segmentation and more complex and computationally demanding workflows. Moreover, image-based metrics are widely used in cell biology and constitute a common strategy in assays such as the quantification of autophagic flux using RFP-GFP-tagged LC3^30^.

Despite these limitations, we have demonstrated that FAβ-gal constitutes a robust, reproducible, fast and unbiased method for quantifying SA-β-gal activity. In this sense, it should be noted that during the writing of this manuscript it came to our knowledge that a similar but more complex procedure was used to evaluate the effect of different senescence modulators, reinforcing the applicability of our approach^30^. Hence, our method overcomes the main limitations of the classical SA-β-gal method with almost no technical changes, thus, having the potential to become a standard procedure in senescence quantification.

## Conclusion

Modern and sophisticated methods to measure cellular senescence have not replaced the classical SA-β-gal assay. This hegemony stems not only from the simplicity, low cost, fast execution and versatility of this method but also from reluctance of researchers to adopt completely new techniques that in most cases are complex. Against this background, we have developed FAβ-gal, a method that combines the advantages of both the original SA-β-gal assay and the fluorescence-based methods for quantifying cellular senescence. This method can be easily introduced into the routine of laboratories currently using the classical colorimetric assay, enhancing the accuracy, sensitivity and reproducibility of the generated results. Thus, FAβ-gal could become a reference method for the routine quantification of cellular senescence as well as a valuable tool for high-throughput analyses such as senescence screenings.

## Material and methods

### Cell culture, buffers, and reagents

GM00038 human fibroblasts were obtained from the Coriell Institute (Camden, NJ) and U20S cancer cells from the American Type Culture Collection (ATCC). GM0038 human fibroblasts and U2OS tumor cells were cultured using Dulbecco’s Modified Eagle Medium (DMEM, Gibco, 41965062) supplemented with 8.7% (v/v) Fetal Bovine Serum (FBS, Corning, 15377636) and 0.87% (v/v) 100x Sodium Pyruvate (Gibco, 11360070), 100x Non-Essential Amino Acids (MEM NEA, Gibco, 11140050), 100x HEPES (Gibco, 15630056), 100× Antibiotic-Antimycotic (Gibco, 15240062) and 100× Penicillin–streptomycin-glutamine (Gibco, 10378016). GM00038 cells were carried to replicative senescence through continuous passage. For SA β-gal assays cells were pooled and counted using a Neubauer chamber and Trypan blue solution (Sigma-Aldrich, T8154). Then, 48 h before SA-β-gal staining, approximately 4000 GM00038 cells/well were seeded in 96-well microscopy plates (Falcon, 353219) previously coated with Poly-L-lysine solution (Sigma-Aldrich, P4707). On the other hand, approximately 2500 U2OS cells/well were seeded in 96-well microscopy plates and, next day, treated with different doses of doxorubicin (Santa Cruz, sc-200923) for 24 h (pulse treatment) or continuously (continuous treatment) to trigger cellular senescence induced by DNA damage. To avoid over confluence, the proliferating control cells of the U2OS doxorubicin experiment were seeded (1500 cells/well) in the 96-well plates 2 days before each time point.

### Fixation and SA-β-gal staining

Cell cultures were fixed for 4 min at room temperature using 4% formaldehyde (Sigma-Aldrich) in phosphate-buffered saline (DPBS, Gibco, 14200083) supplemented with 0.9 mM CaCl_2_ (Merck, 23811000) and 0.49 mM MgCl_2_ (Merck, 1058331000). Immediately, cells were washed 3 times with PBS and then incubated with the staining solution. This solution was prepared following a similar protocol to the one from Itahana et al.^24^: 1 mg/mL 5-bromo-4-chloro-3-indolyl-beta-d-galactopyranoside (X-gal, Thermo Scientific, 10365410), citric acid (7.4 mM) /sodium phosphate (25.3 mM) buffer (pH 6.0), 5 mM potassium ferricyanide (Sigma-Aldrich, 60299) and 5 mM potassium ferrocyanide (Sigma-Aldrich, 60279). We did not include magnesium as proposed in Itahana et al, since mammalian beta-galactosidases do not require metals for the catalysis, as opposed to bacterial beta-galactosidases^31,32^. First, all components of the solution but X-gal were mixed, and pH was adjusted to 5.5. This is critical for the correct functioning of the assay, and its optimal value could vary between 5 and 6, depending on the cell type^24^. Then, X-gal was dissolved in N,N-Dimethylformamide (DMF, Sigma-Aldrich, D4551) at a 20x concentration of 20 mg/mL and added to the prepared solution to obtain the final concentration of 1 mg/mL. Before adding the staining solution, cells were washed with the adjusted solution without X-gal to prevent pH shifts due to PBS remnants. Once the staining solution was added, samples were incubated at 37 ºC under ambient CO_2_ levels. The incubation lasted 24 h in the linearity and tissue assays, and for the specific durations indicated for each time point in the sensitivity assay. After this incubation, cells were checked under the microscope to verify the correct staining and then washed with PBS and stored at 4 ºC until image acquisition.

### Animal experiments

Tissues for histology were obtained from animals that were sacrificed for other experimental purposes under the guidelines of the Committee for Animal Experimentation of the Universidad de Oviedo and the Regional Ethics Committee for Animal Experimentation (PROAE 02/2023). Histology was carried out following a similar protocol to the one of Itahana et al.^24^: tissue samples for histology were rinsed in PBS to remove blood remnants, placed in Tissue-Tek® O.C.T. Compound (Sakura, 4583) and frozen with dry ice. Then, 4 µm sections were cut using a cryostat, placed into superfrost slides and immediately fixed for 1 min at room temperature using 1% formaldehyde. Once fixed, the sections were washed with PBS three times and stained as described in the previous section.

### Image acquisition

Prior to image acquisition, cells were stained with 1 µg/mL Hoechst 33342 (Sigma-Aldrich, B2261) in PBS and, after 10 min in the dark, washed with 0.1% Triton X-100 (Sigma-Aldrich, T9284) in PBS to enhance nuclear staining. Then, after another 10 min in the dark, images were acquired using software Zeiss ZEN 3.3 blue edition in a Zeiss Axio Observer microscope under an A-Plan 10X/0.25 Ph1 objective, using a Colibri 5 LED light source and a halogen lamp for fluorescence and bright field images, respectively. The excitation and emission wavelengths of the four fluorescence channels used — Hoechst, green, red and far-red — are described in Figure S1. Image acquisition was carried out in an automated way from 5 positions around the center of each well using the “Tiles and positions” plugin of Zen software and assisted by software autofocus using Hoechst as the reference channel.

### Image analyses

The generated images were manually curated to eliminate those with debris that could interfere with nuclei count or indigo fluorescence quantification. These clean images were used as input for an analysis pipeline (Figure 2) that quantifies the fluorescence signal and counts the nuclei using BiaPy software^23^. Specifically, the pipeline reads the multichannel TIFF files and calculates the sum of the intensity values (raw integrated density, RawIntDen) of the positive area in the far-red channel. This area is defined by a threshold value set manually using an unstained control image that allows to correct for autofluorescence, i.e., the chosen threshold should be such that the percentage of thresholded area in negative control is very close to 0. Then, it applies an optional rolling-ball background subtraction algorithm to the nuclei channel to enhance subsequent segmentation, and saves the result as a TIFF image that is used as input for BiaPy software, which determines the number of nuclei of each image. Finally, these data are used to compute the CTF/nuclei for each biological replicate (in this case each well), measuring a background area (with no cells or debris) for each experiment to correct background signal.

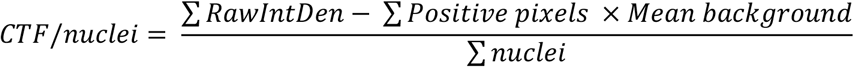

*RawIntDen = Raw integrated density, the sum of the intensity values of the positive pixels Mean background = the mean intensity of an area with no cells*

On the other hand, tissue images were cropped to obtain comparable regions and used as input for this pipeline but skipping the nuclei count step. Thus, in this case we calculated CTF/area (in square micrometers), also correcting background signal.

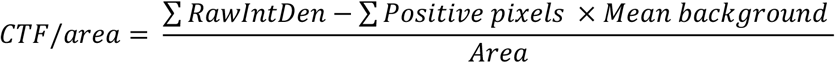

### Autofluorescence analyses

The autofluorescence comparison among channels was conducted with an ImageJ macro (see **Availability and implementation**) that generates a mask of the cells using the green channel and uses it to measure the mean intensity in the green, red and far-red channels. These data were used to calculate the Corrected Mean Cell Fluorescence (CMCF) for each cell to be able to compare among channels and cells. This CMCF was calculated as the mean intensity for each cell and channel after subtracting background fluorescence. On the other hand, the autofluorescence comparison between senescent and proliferative GM00038 human fibroblasts in the different channels was conducted setting a threshold to distinguish background fluorescence from autofluorescence and normalizing the signal by the number of nuclei.

### Experimental design and statistics

All figures in this work correspond to single experiments with the number of biological and technical replicates indicated in the figure legends. All statistical tests and plots were generated using R and RStudio (Core Team, Vienna, Austria, https://www.r-project.org; RStudio Team, Boston, MA, USA, https://www.rstudio.com). In boxplots, the horizontal line represents the median, while the lower and upper limits of the box correspond to the first and third quartiles, and the lower and upper whiskers extend from those limits to the largest/smallest values no further than 1.5 * IQR. (inter-quartile range, i.e. the distance between the first and third quartile). Statistical significance was determined by Student’s unpaired two-tailed t-test, with independent variances and assuming normality, except in the autofluorescence comparison experiment, where a Wilcoxon unpaired two-sample test was used. For correlation studies, a Pearson correlation test was used. The data used to build Supplementary Figure 1 was obtained from FPbase^33^.

### Software

FAβ-gal application was developed using R Shiny. Specifically, RBioFormats package was used for importing images and EBImage^34^ for further manipulations. On the other hand, the Python computational pipeline was developed using BioIO for importing and manipulating the images and BiaPy^23^ together with BioImage Model Zoo^35^ for nuclei segmentation. Moreover, ImageJ^36^ and scikit-image^37^ subtract background algorithms were used in the app and the computational pipeline respectively.

## Code availability and implementation

The source code of FAβ-gal, the R Shiny app, as well as the ImageJ macro for autofluorescence determination are available at [FAβ-gal GitHub repository](https://github.com/GenCanAgeLab/FAB-gal).

## Acknowledgements

We thank Lucas Moledo-Nodar, Víctor Celemín-Capaldi, Pedro M. Quirós and Víctor Quesada for helpful comments and advice. We thank Daniel Franco-Barranco for his assistance in integrating BiaPy nuclei segmentation. This work was supported by the Ministerio de Ciencia, Innovación y Universidades (Spain) (PID2023-148089OB-I00) and Consejería de Ciencia, Innovación y Universidad del Gobierno del Principado de Asturias (AYUD/2021/51062 and IDE/2024/000784). The IUOPA is funded by the Asturian Government and Fundación Cajastur. A.G.T. is supported by a FPU fellowship from Ministerio de Ciencia, Innovación y Universidades (MCIU-24-FPU23-02504). A.P.U. is supported by a Ramón y Cajal grant from Spanish Agencia Estatal de Investigación (RYC2021-031776-I).

## Author contributions

A.P.U. and J.M.P.F. designed the study. A.G.T. carried out most of the experimental work, with support from A.P.U., G.B. and Y.E.. A.G.T. and A.P.U. carried out the statistical and computational studies. A.G.T., J.M.P.F. and A.P.U. wrote the manuscript with input from all authors. A.G.T., D.R.V and A.P.U. wrote the source code and designed the desktop app. All the authors approved the submitted version of the manuscript.

## Disclosure and competing interests statement

The authors declare that they have no conflict of interest.

## Notes

### Competing Interest Statement

The authors have declared no competing interest.

### Summary of Updates

This new version includes the results of applying our method to senescence induced by DNA damage in tumor cells (U2OS), corrects some minor errata and adds Gabriel Bretones as a coauthor.

