## Supplementary figures for "FAβ-gal: an automated fluorescence-based quantification of the senescence-associated beta-galactosidase X-gal assay"

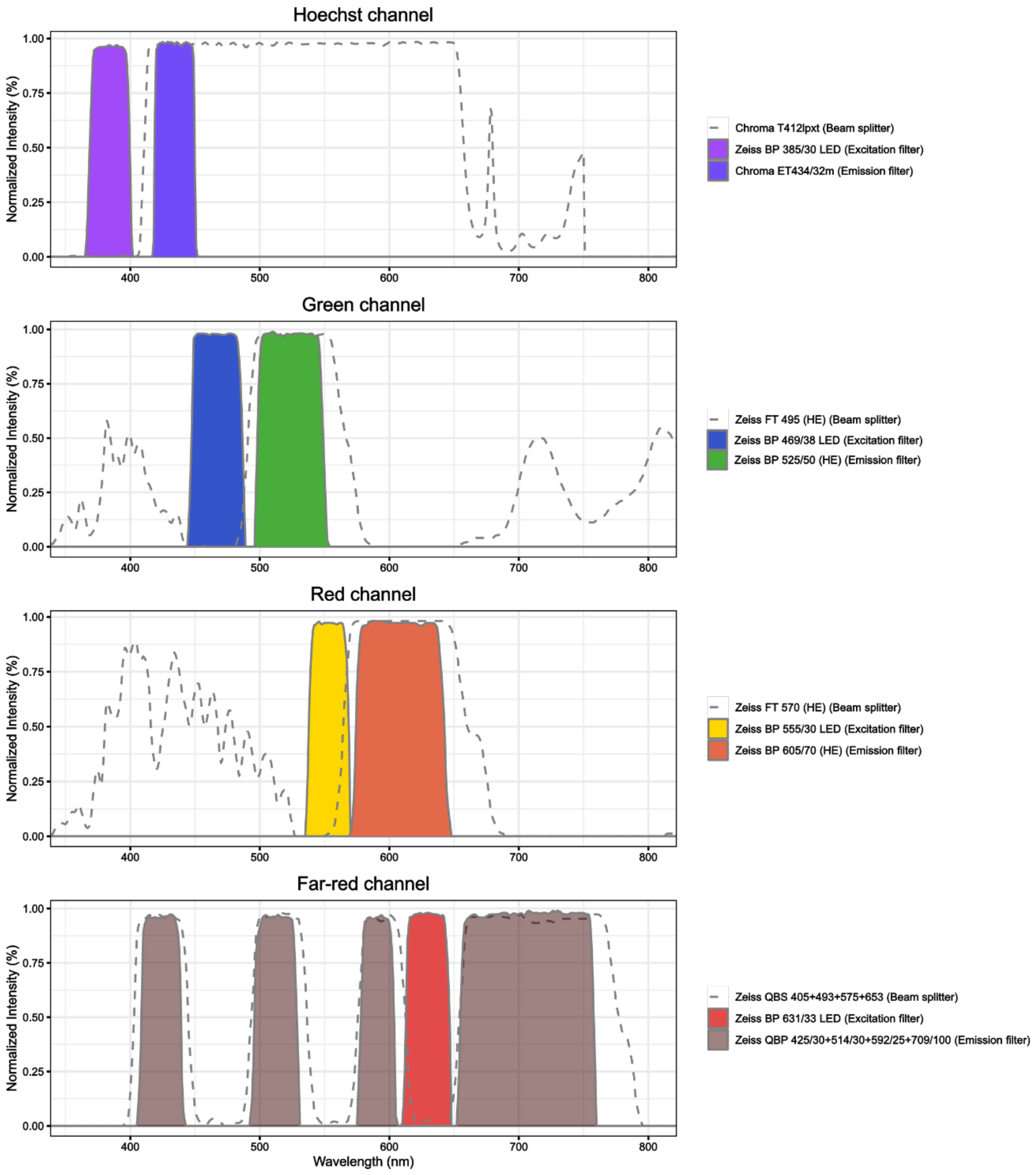


**Figure S1. Microscope configuration for the different channels**

Configuration of the Zeiss Axio Observer microscope for the different channels — Hoechst, Green, Red, Far-red — used in this work. Each plot illustrates the percentage of transmitted light in the displayed electromagnetic spectrum of the different excitation and emission filters and the beam splitters used in each setup.

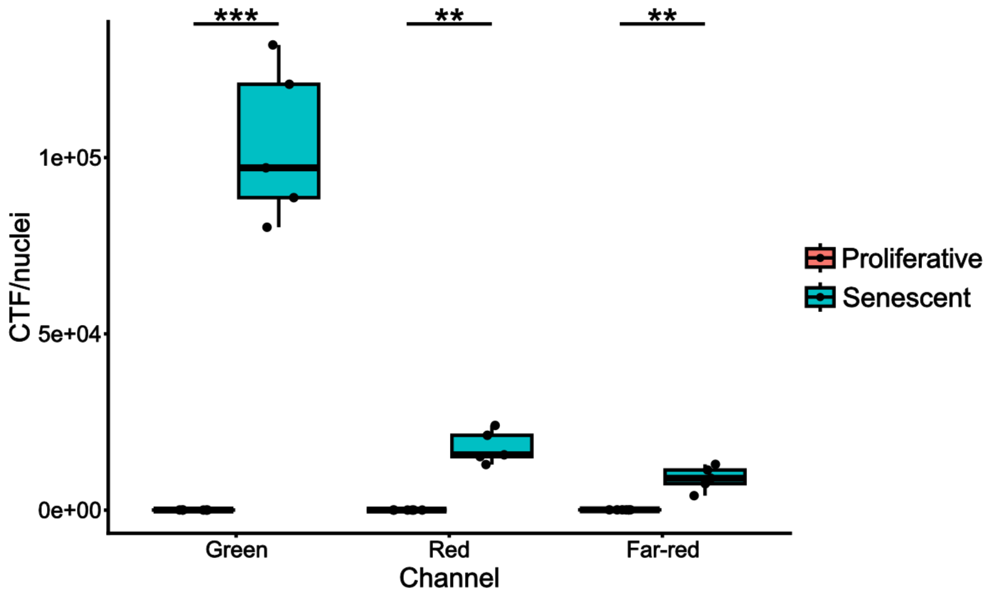


**Figure S2. Autofluorescence is greater in senescent cells than in their proliferative counterparts**

The boxplot represents the Corrected Total fluorescence per nucleus (CTF/nuclei) in the green, red and far-red channels of unstained proliferative and senescent GM00038 human fibroblasts. This CTF was calculated using a threshold to identify autofluorescence, correcting background fluorescence and normalizing by the number of cells. Each point represents the data of a well, which is composed of several images. The statistical significance of the CTF/nuclei differences was calculated using a Student's unpaired two-tailed t-test, with independent variances: **, p-value < 0.01; ***, p-value < 0.001

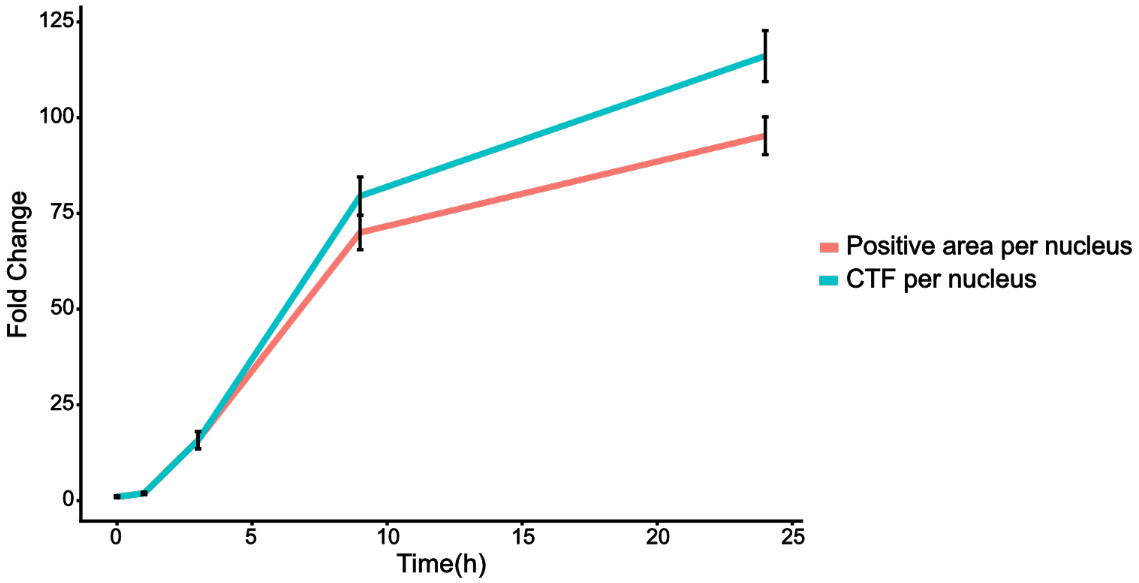


**Figure S3. The CTF per nucleus constitutes a more sensitive metric than the positive area per nucleus**

The line plot represents the relative increase of the Corrected Total Fluorescence per nucleus (CTF per nucleus) and the positive area per nucleus of indigo with assay incubation time in senescent GM00038 human fibroblasts with respect to an unstained control (time 0 h). Error bars represent the standard deviation of the mean for each time point and condition.
